# Selectivity profiles and substrate recognition of Rab phosphorylating kinases

**DOI:** 10.1101/2025.04.09.647999

**Authors:** Deep Chatterjee, Verena Dederer, Landon Vu Nguyen, Marcel Wendel, Kamal R Abdul Azeez, Swetha Mahesula, Florian Stengel, Samara Reck-Peterson, Sebastian Mathea

## Abstract

The Rab GTPase switch-2 region is a hotspot for post-translational modifications. Its phosphorylation can determine whether individuals develop Parkinson’s disease or not. Other modifications of the same region are catalysed by enzymes from bacterial pathogens when they infect human cells. Here, we profiled a set of kinases including LRRK1, LRRK2, DYRK1A, MST1, and TBK1 for their capability of phosphorylating Rab GTPases. We identified several novel kinase:Rab pairs, such as LRRK1:Rab43 and TBK1:Rab29. Further, we comprehensively assessed what makes a Rab GTPase a good kinase substrate, considering the Rab nucleotide binding state and the Rab primary sequence. In a systematic mutational study, Rab variants with modulated phosphorylation properties were established, leading to the identification of a LRRK2 recognition patch in the Rab α3 helix. A Glu to Arg exchange in that patch increased the phosphorylation 18-fold indicating that Rabs are suboptimal LRRK2 substrates. Given that this effect is also observed in a cellular model, we propose that our variants will be excellent tools for analysing the physiological function of Rab phosphorylation.

## Introduction

The Rab family of small GTPases comprises about 70 human proteins with conserved sequence and fold (Pylypenko et al. 2018). Besides their N-terminal GTPase domain, Rab proteins bear an unfolded C-terminal tail. Upon translation, they are bound by cytoplasmic Rab escort proteins (REPs). In the REP:Rab complex, the C-terminal Rab tail is prenylated, allowing for its insertion into membranes. Specific Rabs are associated with specific membranes, thereby acting as labels that assign identity to the membranes, as described for mitochondria (with Rab32 associated), lysosomes (with Rab7A), or the *trans*-Golgi network (with Rab8A) (Homma et al. 2021). Like other GTPases, binding of either GDP or GTP switches the Rabs between distinct conformations. The regions with the largest conformational changes are referred to as the switch-1 and switch-2 regions. Regulators of the guanine nucleotide binding state are GTPase activating proteins (GAPs) that increase GTP hydrolysis thus promoting Rab:GDP complexes, and guanine nucleotide exchange factors (GEFs) that induce GDP release and the subsequent binding of another GTP molecule thus promoting Rab:GTP complexes. Membrane-associated Rab:GTP complexes form anchor points for effector proteins regulating the membrane composition, enabling vesicle budding and fusion, or linking the membrane to motor proteins (Pylypenko et al. 2018).

The Rab docking interface for effector proteins is composed of the switch-1, interswitch and switch-2 regions. The plasticity of the interface and its interactions are regulated *via* a conserved phosphosite in the switch-2 region – about 70% of all Rabs bear a serine or threonine residue in the respective position (Homma et al. 2021). A diverse set of kinases has been reported to phosphorylate Rab proteins at this position, namely LRRK2 (Steger et al. 2016), TAK1 (Levin et al. 2016), TBK1 (Heo et al. 2018), LRRK1 (Hanafusa et al. 2019), MST1/3 (Ueda et al. 2020; Vieweg et al. 2020), and DYRK1A (Wang et al. 2023). While the kinases and their Rab substrates are diverse in terms of subcellular localization and cellular function, a common regulatory mechanism can be hypothesized. One compelling possibility would be that switch-2 phosphorylation is the critical step that enables the binding of a new set of effector proteins (Waschbüsch et al. 2020).

The most well-studied kinase to phosphorylate Rabs in their switch-2 region is LRRK2. Mutations in the gene coding for the LRRK2 protein have been identified as Parkinson’s disease (PD) risk factors (Zimprich et al. 2004). The LRRK2 variants with the highest penetrance exhibit increased kinase activity, linking Rab signalling to neurodegeneration. However, the cellular mechanism by which Rab signalling promotes PD is not fully resolved yet (Taymans et al. 2023). To add another level of complexity, LRRK2 not only phosphorylates Rab proteins, but also acts as a Rab effector protein by being recruited to membranes by specific Rabs. For instance, Rab12 binds to the LRRK2 N-terminus and activates the LRRK2 kinase in cells (Dhekne et al. 2023). Another LRRK2-recruiting Rab protein is Rab29. A mutation in the gene coding for Rab29 has been identified as a genetic risk factor for PD (Guo et al. 2014). Initially, Rab29 was suggested to be a substrate for the LRRK2 kinase (Nirujogi et al. 2021), but more recently, it was shown to bind to the LRRK2 N-terminus and to recruit LRRK2 to membranes in a similar fashion as Rab12 (Zhu et al. 2023).

Rabs and Rab post-translational modifications (PTMs) are not only involved in neurodegeneration, but are also key players when certain pathogenic bacteria infect human cells. Examples for such pathogens include *Legionella* and *Salmonella* species that enter their host cells by endocytosis and reside in a vesicle upon entering (Spanò and Galán 2018). They prevent the cell from maturing the vesicle into a lysosome by modifying the Rabs that are attached to the cytoplasmic face of the vesicle. As a consequence, a different set of effector proteins associates with the vesicle, ultimately turning the vesicle into a bacteria-containing vacuole (Spanò and Galán 2018). An growing list of secreted bacterial enzymes covalently modify Rab proteins by attaching diverse chemical structures. Prominent examples include the enzyme AnkX that attaches phosphorylcholine (Mukherjee et al. 2011), DrrA that attaches AMP (Du et al. 2021), and SseK3 that attaches GlcNAc (Meng et al. 2020). Interestingly, the same switch-2 region that is subjected to LRRK2 phosphorylation is a hotspot for Rab modifications catalysed by pathogen enzymes.

It is well established that Rab proteins and their effectors orchestrate cellular membrane trafficking. Dysregulated membrane trafficking is relevant in neurodegeneration and pathogen infection. The underlying mechanisms are difficult to decipher because they also depend on cell type and developmental stage. Especially the upstream signalling cascades regulating Rab protein activities are incompletely understood. Several kinases have been reported to phosphorylate Rab proteins (Homma et al. 2021), however, a systematic assessment of their substrate selectivities is lacking. To gain insight into Rab regulation, we have determined the substrate selectivity profiles of the known Rab phosphorylating kinases. This also included mapping the respective phosphosites. Further, we have performed an exhaustive mutational study to identify residues that define Rab substrate properties. And finally, we have demonstrated that our Rab variants are differentially phosphorylated in a cellular context as well.

## Methods

### A panel of recombinant Rab proteins

Plasmids for Rab protein expression were purchased from Genscript. They were based upon the pET-28a vector backbone and comprised the respective Rab sequence without the C-terminal prenylation region and a TEV protease-cleavable N-terminal His_6_ tag (see **Table S1** for all insert sequences). For protein expression, the respective plasmid was transformed into *E. coli* BL21(DE3) competent cells. Liquid cultures were grown shaking at 37°C while keeping selection pressure. Once OD600 reached 0.8, temperature was reduced to 18°C and protein expression was induced by adding 0.5 mM IPTG. After 20 hours of incubation, cells were harvested by centrifugation and the cell pellet stored at -20°C until further processing. For protein purification, the cell pellet was resuspended in lysis buffer (50 mM HEPES pH 7.4, 500 mM NaCl, 20 mM imidazole, 0.5 mM TCEP, 5% glycerol, 20 μM GDP, 5 mM MgCl_2_) and sonicated (5 sec pulse - 10 sec pause, amplitude 37%, total pulse time 10 min). After centrifugation (35,000x g, 1 hour, 4°C), the supernatant was loaded onto pre-equilibrated Ni-NTA agarose (Qiagen). Beads were washed with 200 mL lysis buffer and the His_6_-Rab protein was eluted with lysis buffer supplemented with 300 mM imidazole. The N-terminal tag was cleaved by adding TEV protease overnight while dialyzing at 4°C. TEV protease, the cleaved tag and other contaminants were removed by reverse Ni-NTA. The flowthrough was collected, concentrated and subjected to size-exclusion chromatography (20 mM HEPES pH 7.4, 150 mM NaCl, 0.5 mM TCEP, 5% glycerol, 20 μM GDP, 2.5 mM MgCl_2_) using a S200 16/200 column (GE Healthcare) coupled to an ÄKTA pure chromatography system (Cytiva).

### Kinase and phosphatase preparation

The recombinant kinases DYRK1A and MST1 were produced from *E. coli* according to the protocol described above for the Rab panel. The kinases LRRK1, LRRK2 and TBK1 and the phosphatase PPM1H were produced from insect cells as previously described (Preuss et al. 2022; Chatterjee 2022). The respective vectors, boundaries, sequences and tags are listed in **Table S2**.

### Mass spectrometry-based kinase activity assay

The kinases were screened for their ability to phosphorylate individual Rab proteins by applying a mass spectrometry (MS)-based activity assay. The kinases (50 nM) were mixed with Rab substrate (5 μM) in assay buffer (20 mM HEPES pH 7.4, 150 mM NaCl, 0.5 mM TCEP, 1 mM GDP or GTP, 2.5 mM MgCl_2_, and 5% glycerol). The reaction was initiated at 20°C by adding 1 mM ATP and stopped after 5-60 min incubation (depending on kinase activity) by adding MS buffer (0.1% formic acid in water). The samples were analyzed with an Agilent 6230 electrospray ionization time-of-flight mass spectrometer coupled with a liquid chromatography unit 1260 Infinity. 5 μL of the reaction mixture were injected onto a C3 column and eluted at 0.4 ml/min flow rate using a solvent gradient of water to acetonitrile with 0.1% formic acid. Data were acquired using the MassHunter LC/MS Data Acquisition software and analyzed using the BioConfirm vB.08.00 tool (Agilent Technology). The peak intensities of the unphosphorylated and phosphorylated Rab proteins were quantified and displayed with GraphPad Prism.

### Mapping of phosphosites

50 μL of the reaction mixture containing 5 μM phosphorylated Rab protein were freeze-dried in liquid nitrogen before samples were resuspended in 100 μL 8 M urea, reduced with 5 mM TCEP for 30 min at 37°C and alkylated with 10 mM iodoacetamide for 30 min at room temperature. After dilution to 4 M urea with 50 mM NH_4_HCO_3_, a Trypsin/Lys-C mix (Promega) at a protease:protein ratio of 1:25 (w/w) was added, and proteins were digested for 4 hours at 37°C. The solution was diluted with 50 mM NH_4_HCO_3_ to a final urea concentration of 1 M, and proteins were further digested for 16 hours at 37°C. Samples were freeze-dried and kept at -20°C. Prior to MS measurement, tryptic peptides were dissolved in 50 μL 0.1% TFA in H_2_O, acidified with 10% TFA and desalted using PierceTM C18 spin tips (Thermo Fisher). Peptide samples were analyzed on an Orbitrap Fusion Tribrid mass spectrometer (Thermo Scientific) and interfaced with an Easy-nLC 1200 nanoflow liquid chromatography system (Thermo Scientific). Samples were reconstituted in 0.1% formic acid in 5% acetonitrile and loaded onto the C18 analytical column (50 μm × 15 cm, Thermo Scientific). Peptides were resolved at a flow rate of 300 nL/min using a linear gradient of 6-44% solvent B (0.1% formic acid in 80% acetonitrile) over 75 min. Data-dependent acquisition with full scans over a 300−1850 m/z range was carried out using the Orbitrap mass analyzer at a mass resolution of 120,000 at normal mass range, custom automatic gain control target value, and an automatic injection time. The 10 most intense precursor ions were selected for further fragmentation. Only peptides with charge states 2-7 were used, and dynamic exclusion was set to 60 sec. Precursor ions were fragmented using higher-energy collision dissociation (HCD) with normalized collision energy (NCE) set to 30%. An intensity threshold of 2.5e4 was set with an isolation window of 1.6 m/z. Fragment ion spectra were recorded at a resolution of 30,000, automatic gain control target value was set on standard, and the injection time was dynamic. Each sample was measured in technical duplicates. The raw files from LC-MS/MS measurements were analyzed using MaxQuant (version 2.1.4.0) with the following settings: phospho (STY) was set as variable modification, minimum peptide length to 3, maximum peptide length to 35 and match between runs and label-free quantification (LFQ) were enabled. For phosphosite identification data was searched against the specific Rab:kinase pair and processed using Perseus (version 2.0.5.0). Only potential phosphorylation sites with a relative intensity (MS1) of > 10% were taken into account **(Figure S1)**.

### Dynamic scanning fluorimetry (DSF) assay

The assay was performed according to a previously established protocol (Preuss et al. 2022). The Rab protein (4 μM) in assay buffer (20 mM HEPES pH 7.4, 150 mM NaCl, 0.5 mM TCEP, 2.5 mM MgCl_2_, 5% glycerol) was mixed 1:1000 with a SYPRO Orange solution (Sigma-Aldrich). 20 μL of the mixture was pipetted to a 96 well plate and complemented with 1 mM of the nucleotide to be tested (GDP, GTP or GTP-γ-S). The plate was heated gradually from 25° to 95°C and the fluorescence of each sample was monitored using an MX3005P real-time PCR instrument (Stratagene) with excitation and emission filters set to 465 and 590 nm, respectively. The data were analyzed with the MxPro software.

### Dephosphorylation assay

For dephosphorylation of Rab proteins by PPM1H, the pRab (5 μM) was mixed with PPM1H (5 nM) in assay buffer (20 mM HEPES pH 7.4, 150 mM NaCl, 0.5 mM TCEP, 20 μM GDP, 2.5 mM MgCl_2_, and 5% glycerol). The samples were incubated at 20°C for 60 min and analyzed applying the MS method as described for the kinase activity assay.

### Generation of Rab8A variants

Expression plasmids for 92 Rab8A variants were obtained from Genscript (**Table S3**). The variants were expressed and purified as described above for the wild-type protein. For the initial phosphorylation assay, the expression scale was reduced to 50 mL growth medium. For confirmation and quantification of the Rab8A variant substrate properties, the scale was increased to 2 L growth medium.

### Quantification of Rab phosphogrades in cells

All mammalian expression constructs were obtained by subcloning from the bacterial expression constructs into pcDNA3.1(+) *via* KpnI and BamHI cloning sites yielding a fusion protein with an N-terminal eGFP. The plasmid 3xFLAG-LRRK2 was obtained by subcloning LRRK2 from pDEST53 GFP-LRRK2 (Addgene plasmid #25044) into pcDNA5/FRT/TO backbone via BamHI/NotI followed by subcloning the 3xFLAG tag to the N-terminus of LRRK2 via KpnI/BamHI **(Table S4)**. 293T cells (ATCC) were cultured in DMEM (Corning) containing 10% FBS at 37°C and 5% CO_2_ and seeded at a density of 0.3x 10^6^ cells per well in 6-well plate 24 hours before transfection. 293T cells were co-transfected using PEI (Polysciences) with 2 μg of 3xFLAG-LRRK2 and 1 ug of either wild type, E68R, R79A, R104A, or E108R pcDNA3.1-Rab8A. After 48 hours, the wells were treated with either DMSO or 0.5 μM MLi-2 for 4 hours. Cells were washed with PBS and lysed on ice in 300 μL of cold RIPA buffer (50 mM Tris pH 8.0, 150 mM NaCl, 1% Triton X-100, 0.1% SDS, 0.5% Sodium Deoxycholate, 1 mM DTT, cOmplete EDTA-free protease inhibitor cocktail, PhosSTOP phosphatase inhibitor cocktail). Cell lysates were rotated for 15 min and then cleared by centrifuging (13,000x g, 15 min, 4°C). 1 X NuPAGE™ Sample Reducing Agent and 1 X NuPAGE™ LDS Sample Buffer was added to the supernatants and samples were boiled at 95°C for 10 min. For Western blots, 15 μL of the samples were run on a NuPAGE™ 4 to 12 % Bis-Tris gel (Thermo Scientific) for 70 min at 140 V and transferred onto a PVDF transfer membrane (0.45 μm pore size, Thermo Scientific) for 240 min at 200 mA at 4°C. The membrane was blocked in 5% milk powder in TBS-T (20 mM Tris pH 8.0, 150 mM NaCl, 0.1% Tween-20) for 1 hour. Primary antibodies: monoclonal rabbit anti-LRRK2 antibody (Abcam ab133474, 1:1000), monoclonal mouse anti-GFP antibody (Santa Cruz Biotechnology sc-9406, 1:2500), monoclonal rabbit anti-phospho T72 Rab8A antibody (Abcam 230260, 1:1000), or monoclonal rabbit anti-GAPDH antibody (Cell Signalling Technology 14C10, 1:3000) diluted in 1% milk powder in TBS-T, were incubated with the membrane overnight at 4°C. The next day, membranes were washed three times with TBS-T and incubated with IRDye 680RD donkey anti-rabbit IgG and IRDye 800CW donkey anti-mouse IgG secondary antibodies diluted 1:5000 with 4% milk powder in TBS-T for 1 hour at room temperature. The membranes were then washed three times with TBS-T and imaged on a Li-Cor Odyssey CLx controlled by Imaging Studio software. For quantification of protein levels, the signal intensity of each protein band was measured using the Analyze Gels tool in ImageJ (version 2.14.0). Experiments were quantified with n=4, and the phospho-T72 GFP-Rab8A signal was normalized to the LRRK2 signal. Comparative statistical analysis of phosphorylation levels between wild-type Rab8A and mutants was performed with a one-way ANOVA and corrected using Tukey’s multiple comparison test in GraphPad Prism (version 10.1.0).

## Results

### Identification of the *in vitro* Rab substrates for LRRK1, LRRK2, DYRK1A, MST1 and TBK1

The human proteome comprises about 70 Rab proteins. We selected a subset of 31 Rab proteins according to their association with human disease and their reported post-translational modifications. We produced these 31 Rab proteins recombinantly as full-length constructs without the C-terminal prenylation region. The resulting Rab panel (**Table S1**) was probed as substrates for the following Rab kinases, which we also expressed recombinantly: LRRK1, LRRK2, DYRK1A, MST1 and TBK1. The turnover was quantified applying electrospray ionization time-of-flight mass spectrometry (ESI-TOF MS), and the phosphosites were confirmed by proteolytic digestion followed by mass spectrometry (**Figures 1** and **S1**).

**Figure 1.**
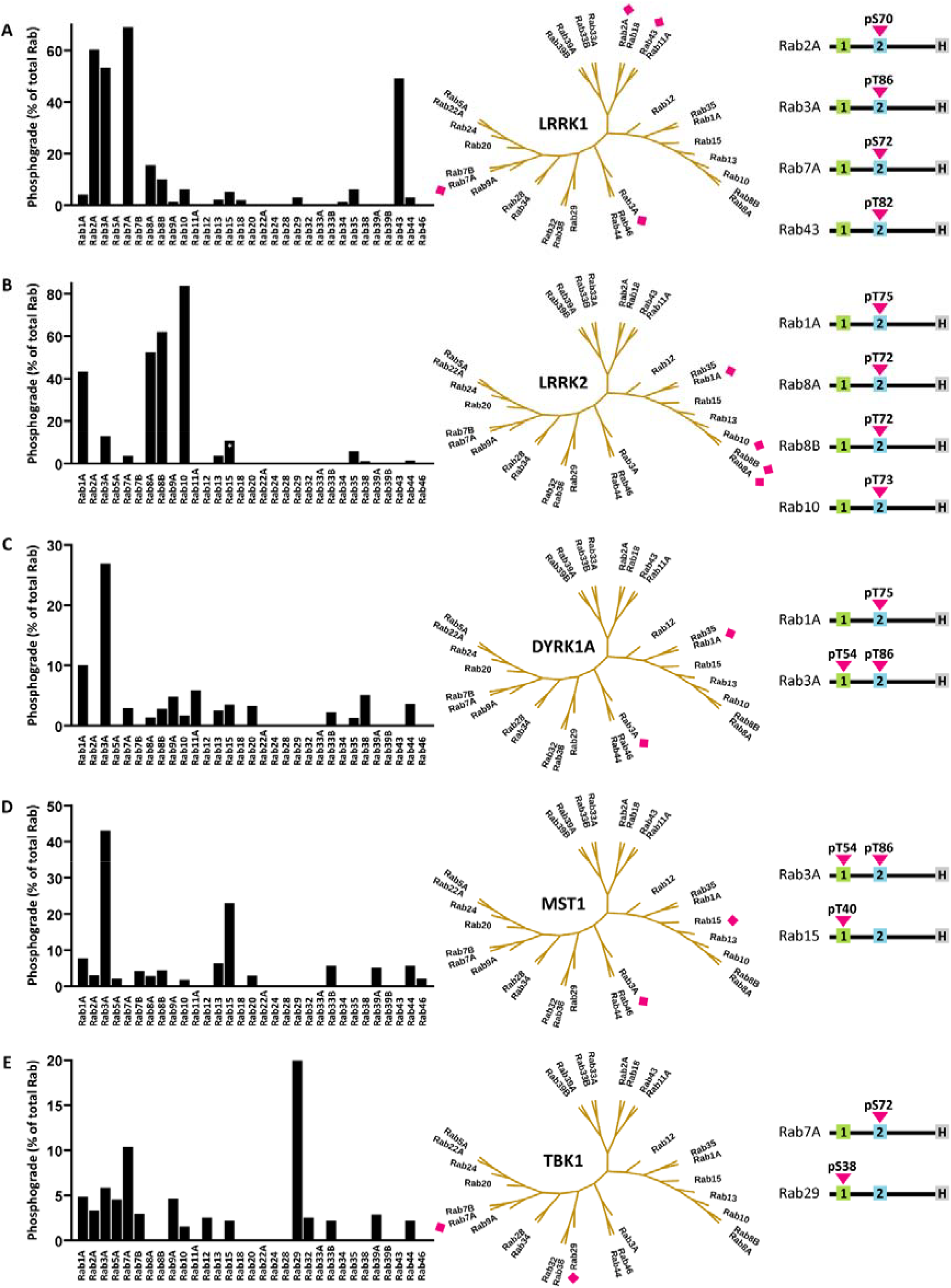
A-E. Probing a panel of recombinant Rab proteins as kinase substrates. The phosphogrades (the percentage of phospho-Rab to total Rab protein) were determined by ESI-TOF MS. Preferred substrates of the respective kinase are labelled with a pink diamond. Further, the phosphosites were mapped using proteolysis and an MS-MS protocol. Most Rabs were phosphorylated in their switch-2 region, but three Rabs were phosphorylated in their switch-1 region. The star in the Rab15 bar indicates that Rab15 was unspecifically phosphorylated at several residues. The detailed results from the phosphomapping are summarized in **Figure S1**.

#### LRRK1

LRRK1 is the most closely related kinase to LRRK2. While gain-of-function mutations in LRRK2 are linked to PD, loss-of-function mutations in LRRK1 are linked to bone disease (Pieridou et al. 2023). Rab7A has been shown to be a LRRK1 substrate (Hanafusa et al. 2019; Malik et al. 2021). Our in vitro assay confirmed this, highlighting the predictive value of our approach. We also identified Rab2A, Rab3A and Rab43 as potential novel LRRK1 substrates. As expected for a stringent substrate recognition mechanism, substrates were phosphorylated exclusively in the conserved Ser/Thr residue in the Rab switch-2 region. Interestingly, the LRRK1 Rab substrates were not closely related, but scattered over the Rab phylogenetic tree (Figures 1A and S1).

#### LRRK2

Due to its association with PD, LRRK2’s substrate specificity has been comprehensively studied. In our assay we identified four Rabs previously described as LRRK2 substrates: Rab1A, Rab8A, Rab8B and Rab10 (Steger et al. 2016). Three Rabs expected to be LRRK2 substates, Rab12, Rab29, and Rab43, were not phosphorylated by LRRK2 in our *in vitro* assay. These Rabs may not be direct LRRK2 substrates, even though their cellular phosphogrades depended on the LRRK2 kinase activity in cell-based assays (Nirujogi et al. 2021). As expected, all LRRK2 substrates were phosphorylated in the conserved Thr residue in the Rab switch-2 region. In contrast to the LRRK1 substrates, the LRRK2 substrates clustered in one branch of the Rab phylogenetic tree (**Figures 1B** and **S1**).

#### DYRK1A

Increased DYRK1A activity is linked to several neurodevelopmental disorders including Down syndrome (Duchon and Herault 2016), but has only recently been proposed as a Rab-phosphorylating kinase (Wang et al. 2023). In our reconstitution assay, Rab1A and Rab3A were identified as the best DYRK1A substrates. Both Rabs were phosphorylated in their switch-2 region (**Figures 1C** and **S1**). DYRK1A is known to have a preference for the linear motif `-RXXS/TP-` (Soundararajan et al. 2013). In line with this preference, both Rab1A and Rab3A bear an arginine residue in position -3 relative to the phosphosite. The physiological significance of DYRK1A phosphorylating Rab proteins in their switch-2 region remains to be investigated.

#### MST1

MST1 has been proposed to orchestrate Rab signalling in lymphocytes (Ueda et al. 2020). When testing our recombinant Rab panel as potential MST1 substrates, Rab3A and Rab15 stood out with the highest phosphogrades. Interestingly, Rab3A was phosphorylated both in the switch-1 and the switch-2 region, while Rab15 was phosphorylated exclusively in the switch-1 region (**Figures 1D** and **S1**). The activity of MST1 towards Rab8A was much lower than towards Rab3A and Rab15. The previously described MST3 phosphosite S111 in Rab8A was not detected, potentially reflecting differences in specificity between MST1 and MST3 (Vieweg et al. 2020).

#### TBK1

TBK1 functions in innate immunity, and mutations in TBK1 have been linked to both amyotrophic lateral sclerosis (ALS) and frontotemporal dementia (FTD) (Harding et al. 2021; McCauley and Baloh 2019). Our assay confirmed that Rab7A is a TBK1 substrate (Heo et al. 2018). As expected, Rab7A was phosphorylated in its switch-2 region. The best TBK1 substrate, however, was Rab29 (**Figure 1E**). None of the other tested kinases had phosphorylated Rab29 indicating that the protein adopted a stable conformation without accessible unfolded regions. In contrast to Rab7A, Rab29 was phosphorylated in its switch-1 region (**Figure S1**). The mechanistical and physiological implications of this phosphorylation remain to be determined.

### Determination of the role of nucleotide binding on Rab protein stability

Next, we investigated the role of the bound guanine nucleotide for all 31 Rabs (**Figure 2A**). We started by comparing the temperature stabilities of apo Rab8A, Rab8A:GDP and Rab8A:GTP. The apo protein displayed the lowest thermostability. GDP stabilized Rab8A dramatically by about 15 K. The presence of GTP stabilized the protein even further by additional 6 K (**Figure 2B**). This is in line with our current understanding of Rab regulation, where GDP is thought to bind less tightly than GTP, thereby facilitating GEF-induced nucleotide exchange (Pylypenko et al. 2018). We then tested the temperature stabilities of our complete Rab panel. Many of the apo Rabs did not display defined melting curves indicating an unstable fold. Due to the high affinity of Rabs for guanine nucleotides and their high concentrations in cells, apo Rabs are not expected to be found in cells (Pylypenko et al. 2018). All Rabs from the panel bound to and were stabilized by guanine nucleotides. The melting temperatures of the Rab:GDP and Rab:GTP complexes were between 45 and 78°C (**Figure 2C**). Interestingly, for several Rabs including Rab9A and Rab13 the Rab:GDP and Rab:GTP complexes displayed equal stability. This suggests that for these Rabs the nucleotide complexes are structurally similar. Importantly, the temperature stability assay also served as a quality control for the Rab panel. The high melting temperatures indicated that the Rabs retained their native fold during the phosphorylation assays. No correlation between Rab thermostability and Rab substrate properties was observed.

**Figure 2.**
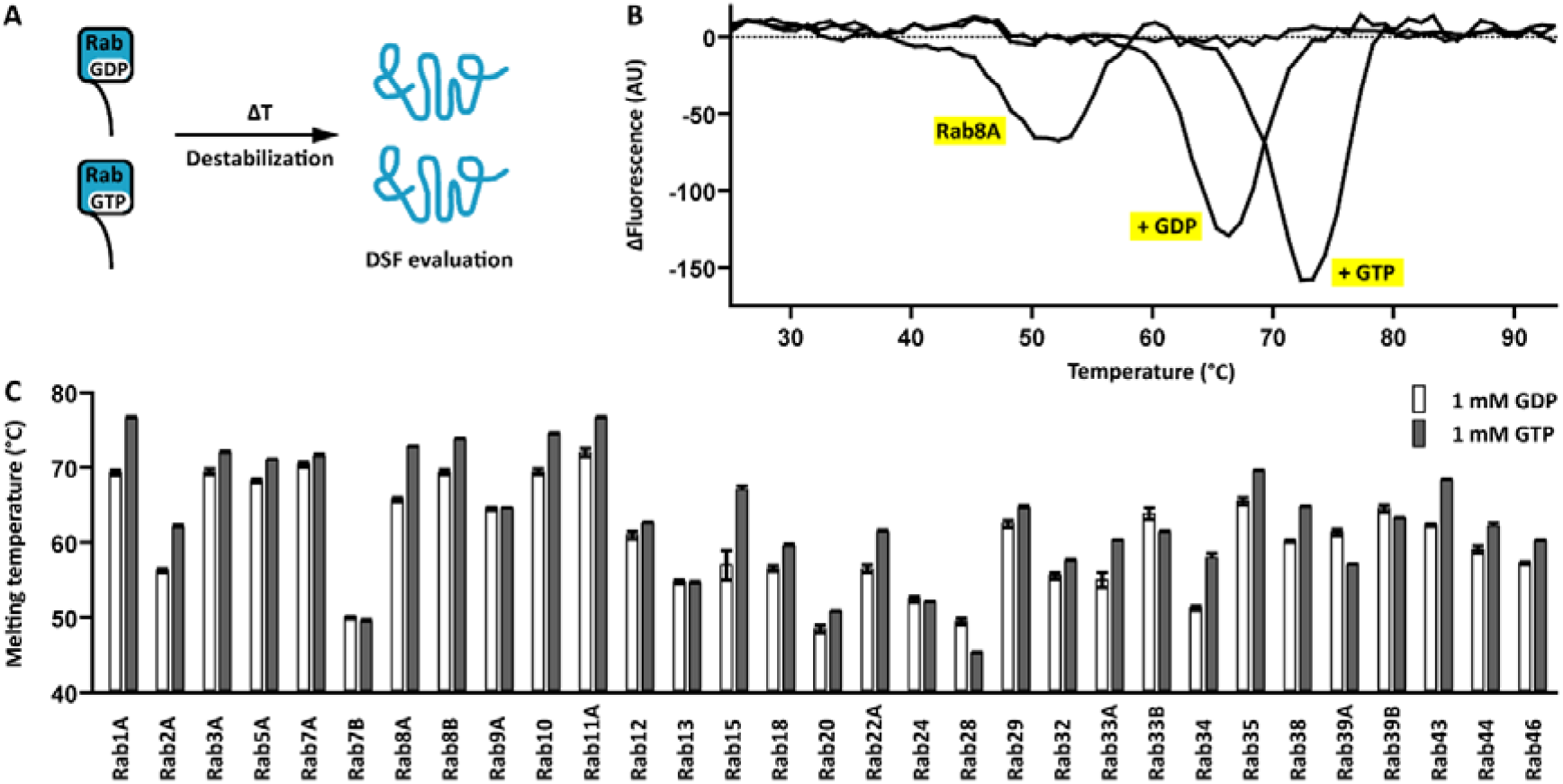
Thermostability of Rab:GDP and Rab:GTP complexes. **A**. Principle of the dynamic scanning fluorimetry (DSF) assay. The temperature of the samples is gradually increased resulting in protein destabilization. Fluorescence is monitored to measure protein unfolding. **B**. Apo Rab8A unfolded at 51°**C**. The addition of GDP and GTP stabilized Rab8A by 15 and 21 K, respectively. C. All Rabs from the panel were stable in presence of GDP and GTP with melting temperatures between 45 and 78°C.

### Rab substrate properties depend on the bound guanine nucleotide

Basically all aspects of Rab biology depend on whether the Rab is bound to GDP or to GTP. Therefore, it is likely that Rab phosphorylation is also regulated by the bound guanine nucleotide. To test this hypothesis, Rab:GDP and Rab:GTP complexes were prepared for eight selected Rabs and the complexes were tested as LRRK1 and LRRK2 substrates (**Figure 3A**). LRRK2 preferentially phosphorylated the Rab:GDP complexes. For example, the Rab10:GDP phosphograde was 4-fold higher than the Rab10:GTP phosphograde. This observation can be explained mechanistically by considering the properties of the GTP γ phosphate, which is situated at the centre of the Rab:GTP complex, coordinating the switch-1 and switch-2 regions and condensing the complex. The γ phosphate would help maintain the switch-2 region in a folded state, making it less accessible for phosphorylation (**Figure 3B**). In the Rab:GDP complex, in contrast, the switch-2 region is more flexible and thus more accessible for phosphorylation. Interestingly, this model was not valid for LRRK1 substrates. In contrast, we found that LRRK1 phosphorylated most Rab:GDP and Rab:GTP complexes at similar rates (**Figure 3A**).

**Figure 3.**
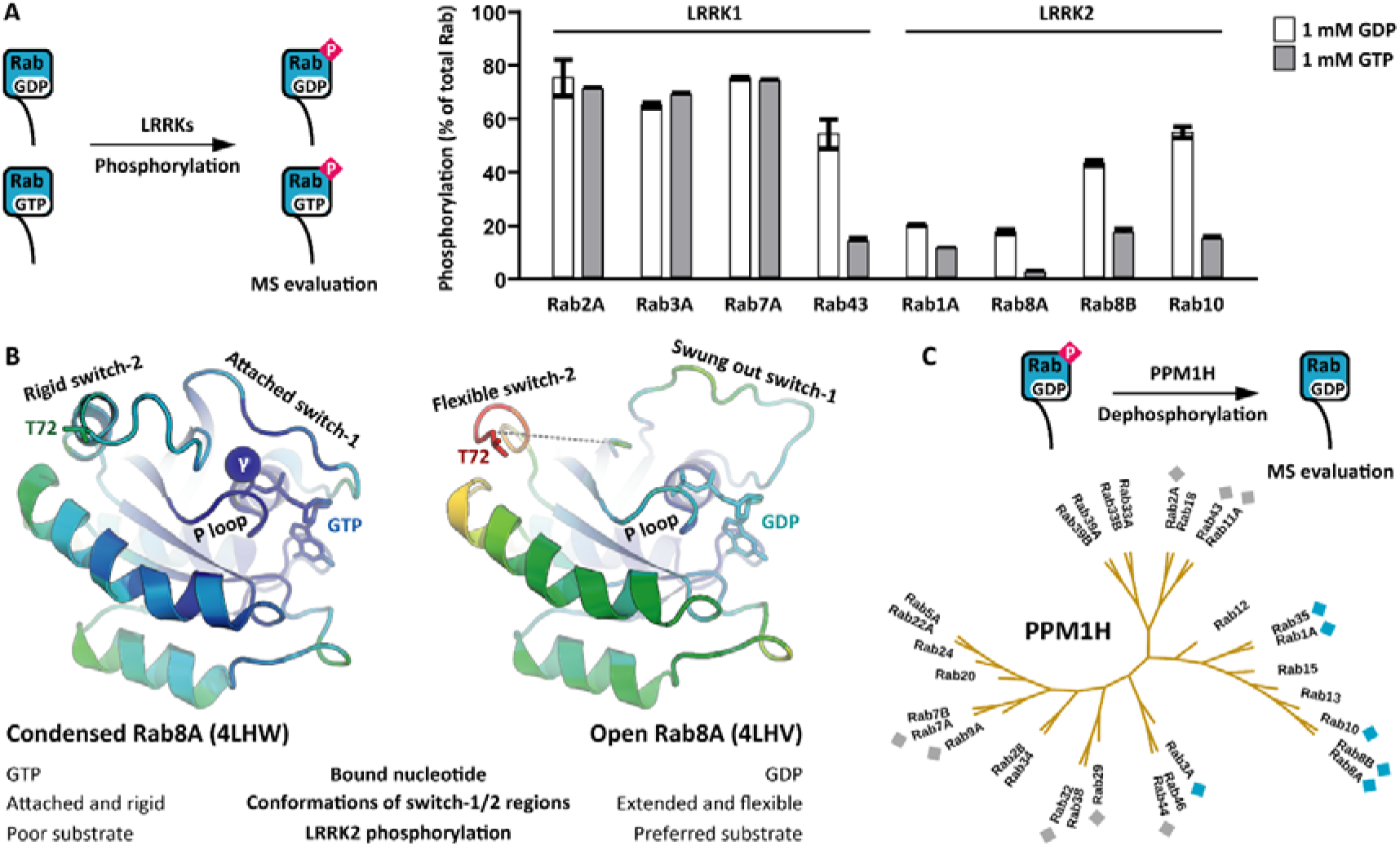
LRRK2 preferentially phosphorylates Rab:GDP complexes. **A**. Selected Rab:GDP and Rab:GTP complexes were subjected to LRRK1 and LRRK2 phosphorylation. In contrast to LRRK1, LRRK2 preferred Rab:GDP complexes as substrates. **B**. Structure models of Rab8A:GDP and Rab8A:GTP highlighting the impact of the GTP γ phosphate on Rab8A substrate properties. Blue indicates low B-factors, while red indicates high B-factors. **C**. pRabs were tested as PPM1H phosphatase substrates by incubating the pRabs in presence of PPM1H followed by MS evaluation of the phosphogrades. The pRabs labelled with blue diamonds were completely dephosphorylated, while the pRabs labelled with grey diamonds were not dephosphorylated by PPM1H.

### Selectivity of the phosphatase PPM1H

The phosphatase PPM1H is known to dephosphorylate pRab proteins (Berndsen et al. 2019). With a panel of pRabs from our reconstitution assay available, we investigated PPM1H substrate specificity. Interestingly, only phosphorylated LRRK2 substrates such as pRab8A and pRab10 were dephosphorylated by PPM1H (**Figure 3C**). The pRabs resulting from phosphorylation with other kinases were not dephosphorylated by PPM1H. Consistent with our finding that Rab29 and Rab43 were not LRRK2 substrates, these Rab were also not dephosphorylated by PPM1H. As reported previously, PPM1H is a true LRRK2 antagonist (Phung et al. 2024), and PPM1H and LRRK2 substrate recognition may rely on the same residues in the Rab sequence (Waschbüsch et al. 2021).

### Mutational analysis of LRRK2 substrate recognition *in vitro*

Next, we sought to identify Rab residues that were important for LRRK2 substrate recognition. The *bona fide* LRRK2 substrate Rab8A was chosen as a model substrate because of its favourable expression and purification properties. In the enzyme-substrate complex, ATP is bound to the LRRK2 active site. The Rab8A phosphoacceptor residue T72 must be positioned near the ATP γ phosphate. Further, in analogy to many other kinase substrates, the residues flanking T72 likely form a β strand that interacts with the LRRK2 activation segment (Miller and Turk 2018). Taking these geometric restraints into account, there are about 50 Rab8A residues that could potentially interact with the LRRK2 kinase domain in the enzyme-substrate complex. These residues are situated in the Rab8A P loop, in the switch-1 and switch-2 regions, and in the α3 helix. To comprehensively investigate the role of these amino acids in LRRK2 substrate recognition, we performed an alanine scan. Additionally, charged residues were mutated to residues with the opposite charge, and small residues to large residues. Altogether, we generated and expressed 92 Rab8A variants, which were then purified using a one-step purification protocol (**Table S3**). Several variants, including most P loop variants, exhibited solubility issues. The remaining 71 variants were assessed in our reconstitution assay and their phosphogrades relative to wild-type Rab8A were determined. Several Rab8A variants exhibited either increased or decreased T72 phosphogrades. These residues were situated mainly in the switch-2 region or in the C-terminus of the α3 helix (**Table S3**). To confirm these findings we purified to homogeneity all variants with phosphogrades less than 50% or more than 200% in comparison to wild-type Rab8A and then determined their phosphogrades (**Figure 4A**).

**Figure 4.**
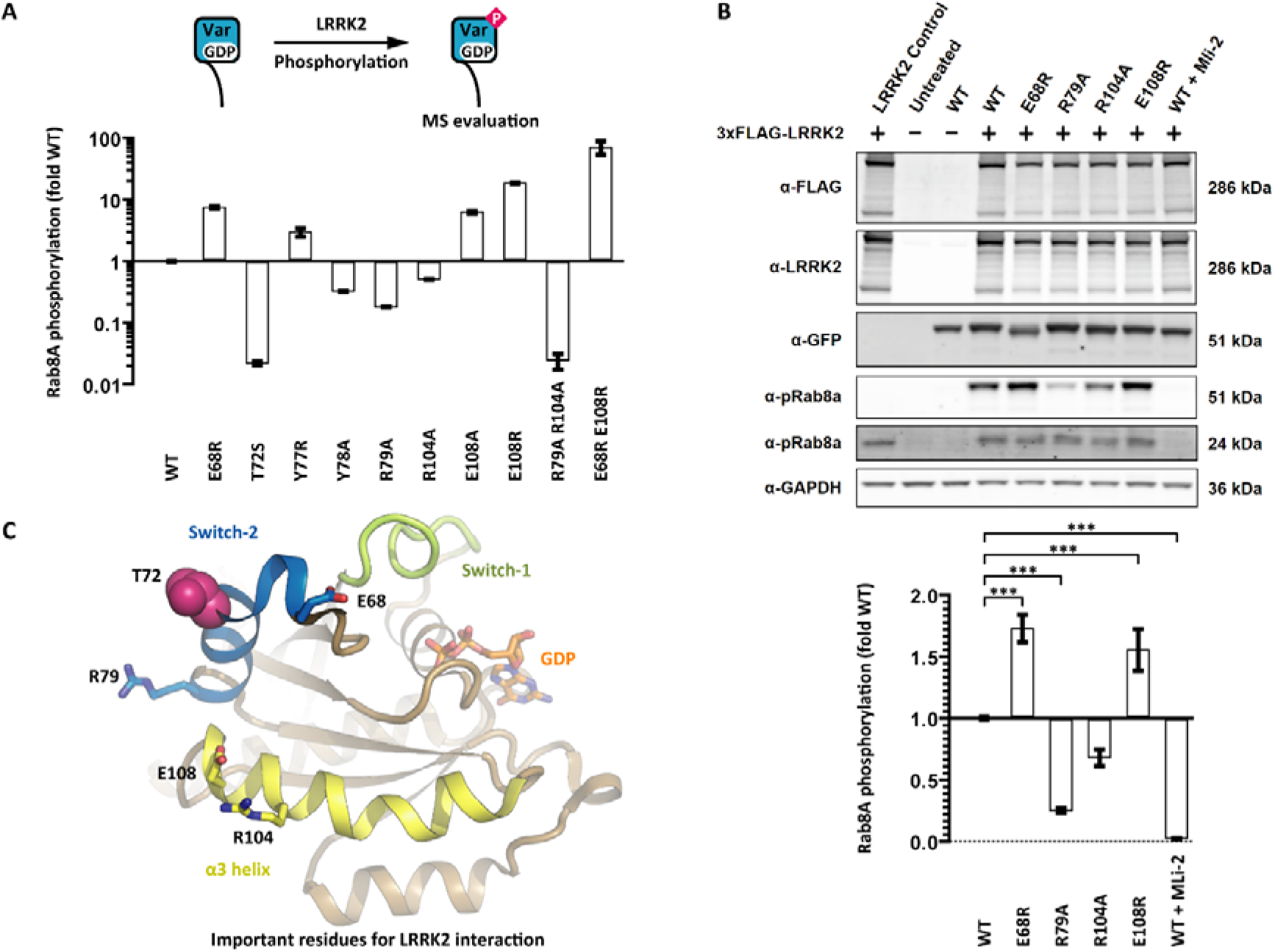
Identification of Rab8A residues that modulate LRRK2 substrate recognition. **A**. 92 Rab8A variants were tested as LRRK2 substrates. Several variants were poor substrates in comparison to wild-type Rab8A. Other variants boosted the LRRK2 kinase activity nearly 100-fold. **B**. Recombinant LRRK2 and eGFP-Rab8A variants were expressed in 293T cells and the Rab8A phosphogrades were assessed by Western blotting. The phosphogrades of the Rab8A variants R79A and R104A were reduced, and the phosphogrades of the variants E68R and E108R increased. The results of at least three independent measurements are shown as mean ± SD, *** indicates P <0.001. **C**. Rab8A:GDP structure model (from PDB ID 4LHY) with the T72 phosphoacceptor residue and residues that modulate the substrate properties for the LRRK2 kinase. Our mutational analysis suggests that the LRRK2 docking interface is in the C-terminus of the α3 helix.

Mutating the phosphoacceptor residue T72 to serine resulted in a variant that was not significantly phosphorylated by LRRK2. The preference of LRRK2 for threonine over serine has also been observed for peptide substrates (Pungaliya et al. 2010). This is in line with the observation that kinases with a small or branched residue in the DFG+1 position (isoleucine in LRRK2) generally prefer threonine residues as phosphoacceptors (Chen et al. 2014). However, this rule of thumb is not valid for LRRK1 (also isoleucine in DFG+1 position). The preferred LRRK1 substrate is Rab7A with the phosphoacceptor residue S72 (**Figure 1A**).

As expected, residues in the Rab8A switch-2 region were also important for LRRK2 substrate recognition. Mutating any residue between the -3 to +3 position relative to the phosphoacceptor led to Rab8A variants with phosphogrades of <50% in comparison to wild-type Rab8A. In contrast, variants of the residues in -4 and +5 positions were better substrates for LRRK2. The phosphogrades of Rab8A E68R and Y77R were increased 7-fold and 3-fold, respectively (**Figure 4A**).

A surprising impact on the Rab8A substrate properties was observed for residues in the C-terminus of the α3 helix. In Rab8A R104A the phosphograde was reduced 2-fold, while E108R led to an 18-fold increase. The α3 helix is distant to the phosphoacceptor. We therefore suggest that the patch formed by R104 and E108 serves as a docking interface for LRRK2 in substrate recognition (**Figure 4C**).

Finally, we combined impactful mutations and assessed the substrate properties of the resulting proteins. We found that the double variant R79A R104A was a 50-fold worse substrate for LRRK2 than wild-type Rab8A. In contrast, the double variant E68R E108R showed an 80-fold increase in phosphograde score (**Figure 4A**). Our results suggest that LRRK2’s Rab substrates have evolved to be suboptimal LRRK2 kinase substrates.

### Mutational analysis of LRRK2 substrate recognition in cells

Next, we turned to a cell-based assay to further investigate the LRRK2 substrate recognition. FLAG-tagged LRRK2 and eGFP-tagged Rab8A variants were co-expressed in 293T cells. Their expression levels and the Rab8A phosphogrades were evaluated by Western blotting (**Figure 4B**). In the absence of recombinant FLAG-LRRK2, no Rab8A phosphorylation was observed. The expression of FLAG-LRRK2 resulted in phosphorylation of both endogenous Rab8A and recombinant eGFP-Rab8A. This phosphorylation was completely abrogated in the presence of the LRRK2 inhibitor MLi-2. The expression levels of all Rab8A variants were identical, however, the apparent molecular weight of the variant Rab8A E68R differed slightly from wild-type Rab8A, potentially indicating differences in post-translational modifications (PTMs). The phosphogrades of all tested Rab8A variants significantly differed from wild-type Rab8A with E68R and E108R being more phosphorylated and R79A and R104A being less phosphorylated. While the magnitudes of the effect were modest compared to our reconstitution assay, the same trends were observed.

## Discussion

The identification of the physiologic substrates of a given kinase remains challenging. Here, we used *in vitro* reconstitution assays to profile five distinct Rab kinases. The potential substrates from the Rab panel were near full-length proteins in their native conformation, bound to their physiologic co-factor. Therefore, we were able to address several levels of substrate recognition, including the phosphoacceptor residue, its flanking sequence and potential docking interfaces distal to the phosphoacceptor. All kinases were catalytically active and phosphorylated a subset of Rabs. Moreover, their selectivity profiles were unique, indicating that switch-2 phosphorylation is a common mechanism beyond LRRK2 and its known substrates. All established kinase:Rab pairs including LRRK1:Rab7A (Hanafusa et al. 2019), LRRK2:Rab10 (Steger et al. 2016) and TBK1:Rab7A (Heo et al. 2018) were detected in our reconstitution assay highlighting its predictive potential. Several Rabs discussed as LRRK2 substrates previously (Nirujogi et al. 2021) were not positive in our reconstitution assay, possibly because these Rabs (Rab12, Rab29 and Rab43) are not direct LRRK2 substrates, but rather are important for recruiting LRRK2 to membranes (Dhekne et al. 2023; Zhu et al. 2023). We also discovered new kinase:Rab pairs, showing that LRRK1 phosphorylates Rab43 in its switch-2 region and TBK1 phosphorylates Rab29 in its switch-1 region.

Rab protein variants with modulated mechanistic properties are valuable tools for the investigation of cellular Rab functions. To expand the existing toolkit, we tested 92 Rab8A variants in our reconstitution assay aiming to identify variants with modulated substrate properties. In addition to confirming the significance of the phosphoacceptor residue and its flanking sequence, a potential docking interface for LRRK2 distal to the phosphoacceptor was discovered. Mutating residues in the C-terminus of the Rab8A α3 helix modulated phosphorylation by LRRK2. Single variants displayed activities between 0.5-fold and 18-fold in comparison to wild-type Rab8A. It is important to note that the identified residues do not contribute to nucleotide binding and GTP hydrolysis. Interestingly, the same docking interface is involved in PPM1H substrate recognition. Here, the C-terminus of the α3 helix interacts with the flap region of PPM1H (Waschbüsch et al. 2021). This also explains our finding that PPM1H dephosphorylated exclusively LRRK2 substrates (**Figure 3C**). The same docking interface in the C-terminus of the Rab α3 helix mediates substrate recognition of both LRRK2 and PPM1H.

Recombinant LRRK2 constructs display low catalytic activities with turnover numbers in the range of 0.002 s^-1^. This is the case for both the full-length protein and truncated forms. One explanation for the low catalytic activity is that LRRK2 adopts an autoinhibited conformation and that the observed activity is due to ‘leaky autoinhibition’. For the homologous kinase LRRK1, a regulatory mechanism has been established. Several isoforms of protein kinase C (PKC) activate LRRK1 by phosphorylating a loop in the CORB region distal to the kinase domain (Malik et al. 2022). In its non-phosphorylated form, LRRK1 is autoinhibited with the CORB loop blocking the kinase active site. Phosphorylation leads to the displacement of the loop and subsequent kinase activation (Reimer et al. 2023). An analogous mechanism for LRRK2 activation remains to be discovered (Taymans et al. 2023). Another explanation for the low catalytic activity is that LRRK2 has evolved to be an inefficient kinase similar to a concept described for cyclin-dependent kinases (CDKs) (Miller and Turk 2018). This hypothesis is supported by the fact that mutating the physiological LRRK2 substrate Rab8A in only two residues boosted the turnover number 80-fold to almost 0.2 s^-1^. Whether the low catalytic activity of recombinant LRRK2 is due to ‘leaky autoinhibition’ or ‘catalytic inefficiency’ needs to be established in further studies.

There are two main possibilities for the physiological significance of the Rab switch-2 phosphorylation. The first is that the phosphorylated switch-2 region enables the binding of a new set of effector proteins, as demonstrated for pRab8A:RILPL2 (Waschbüsch et al. 2020). The second is that switch-2 phosphorylation locks the pRabs in an inactive state. We have demonstrated that LRRK2 preferably phosphorylated Rabs bound to GDP (**Figure 3A**). In the cell, pRab:GDP is differentially regulated in comparison to Rab:GDP due to steric hindrance of protein interactions. For example, Rab GEFs such as RABIN8 and GDI1 do not interact with pRab:GDP. The inactive pRab:GDP complex may remain associated with the membrane ‘on hold’ awaiting dephosphorylation.

The Rab GTPase switch-2 region is a hotspot for PTMs. In addition to the aforementioned cellular kinases, there are numerous enzymes secreted by pathogenic bacteria that modify the switch-2 region in the course of pathogen infection. The pathophysiological relevance of these PTMs is obvious: Finetuning of the phosphorylation grade determines whether an individual develops PD or not. And pathogen-catalysed Rab modifications determine whether a given vesicle turns into a lysosome or into a bacteria-containing vacuole. It can even be hypothesized that cellular kinases and pathogen enzymes spatiotemporally compete for attaching their respective PTM to the Rab switch-2 region. The methods and results presented in this manuscript are the basis for future studies addressing this hypothesis.

---

**Figure S1.**
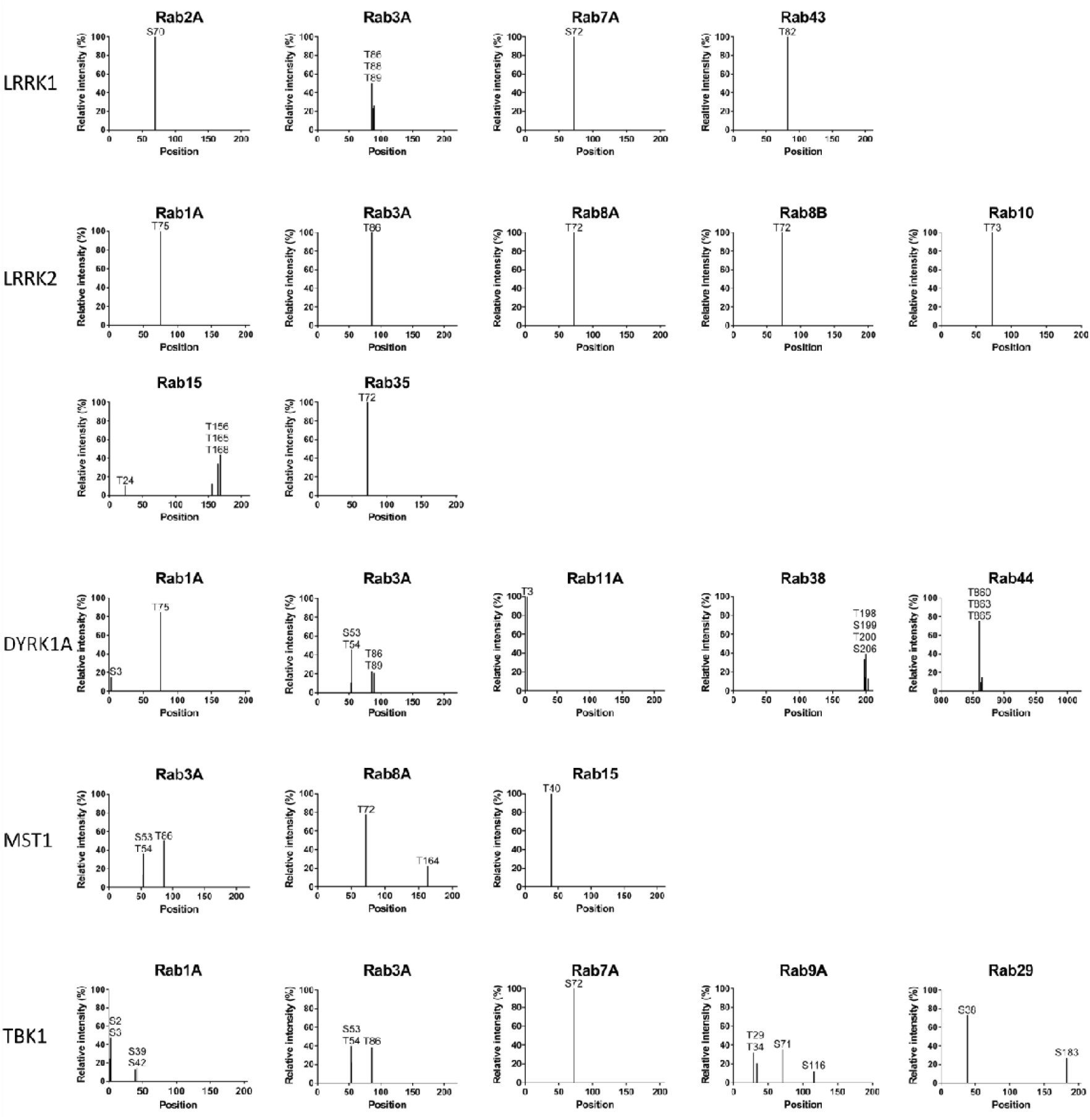
Mapping of phosphosites in Rab proteins. Selected samples from the reconstitution assay (**Figure 1**, left side) were additionally subjected to proteolytic digest and analyzed *via* mass spectrometry for mapping of phosphosites. Bar charts indicate the relative intensity of the detected phosphosites for each kinase:Rab pair. Only phosphorylation sites with intensities >10% are shown.

